# Fungal communities associated with the eastern larch beetle: diversity and variation within developmental stages

**DOI:** 10.1101/220780

**Authors:** Audrey-Anne Durand, Jean-Philippe Buffet, Philippe Constant, Eric Déziel, Claude Guertin

**Author notes:** Corresponding Author: Claude Guertin, 531 Boul. Des Prairies, Laval, Quebec, Canada, H7V 1B7.

## Abstract

Bacterial and fungal communities associated with bark beetles, especially some *Dendroctonus* species, mediate challenging aspects of the subcortical habitat for their hosts. Filamentous fungi and yeasts are important in various metabolic processes of certain bark beetles, with involvement in nutrition, protection and detoxification of plant defense compounds. The eastern larch beetle, *Dendroctonus simplex*, is recognized as a serious forest pest in the upper part of North America. Although *D. simplex* is well studied, the fungal communities and their ecological role remain to be investigated. Here, using 18S rRNA gene pyrosequencing, we provide a comprehensive overview of the yeasts and filamentous fungi associated with the eastern larch beetle and compare fungal communities between different developmental stages and microenvironments. Fungal mycobiome associated with the galleries was also investigated. Our study has unveiled an unexpected fungal diversity associated with the developmental stages. Significant differences in species richness between the developmental stages were determined. Yeasts were found to be predominant in the adult and larval stages, whereas filamentous fungi were most prevalent in the pupae. Our results indicate a possible implication of yeasts in the eastern larch beetle nutrition.

## Introduction

The bark beetle-microbe community is a complex assemblage that has fascinated ecologists and microbiologists for nearly a century. Indeed, bark beetles, especially *Dendroctonus* species (Coleoptera: Curculionidae: Scolytinae), are among the most ecologically important organisms in conifer forest ecosystems worldwide. They colonize the subcortical tissues of conifer with the aid of their microbiota, including filamentous fungi, yeasts and bacteria (Popa *et al*., 2012; Six, 2013). These associated microorganisms play key roles in bark beetles’ biology, including nutrition, protection against pathogens, detoxification of plant defense compounds for host plant use, and swarm behavior modifications by the synthesis of pheromones (Popa *et al*., 2012; Adams *et al*., 2013; Davis, 2014; Shi *et al*., 2014; Hu *et al*., 2015).

*Dendroctonus* species are commonly associated with filamentous fungi and yeasts (Rivera *et al*., 2009; Popa *et al*., 2012; Six, 2012; Davis, 2014; Hofstetter *et al*., 2015). The most prevalent bark beetle fungal partners are Ophiostomatales belonging to the *Grosmannia* and *Ophiostoma* genera, and yeasts of the *Saccharomycetaceae* family (Six *et al*., 2011; Popa *et al*., 2012; Davis, 2014). Bark beetles carry their fungal communities in the gut, on the surface of their exoskeleton or within highly specialized structures (Six, 2003; Six, 2012; Davis, 2014). As bark beetle tunnel under the bark, they inoculate the wall of their galleries with the fungus (Paine *et al*., 1997; Six & Klepzig, 2004).

While most bark beetle are phloeophagous, some are considered mycophloeophagous - gaining nutriments from feeding on fungi as well as phloem (Popa *et al*., 2012; Six, 2013). Some of these nutritional symbioses are obligate where the beetle gains nitrogen and others nutrients from the fungi (Ayres *et al*., 2000; Bleiker & Six, 2007). Feeding on Ophiostamatales fungi increases beetles fitness throughout their developmental stages by stimulating the growth of larvae as well as increasing the fecundity, reproduction and survival rate of adults (Moore & Six, 2015).

Yeasts, known to be prolific metabolizers, also have significant functional roles in bark beetle ecology. Some beetle-associated yeasts use terpene defenses as carbon sources (Davis, 2014). Furthermore, yeasts isolated from several *Dendroctonus* species produce a variety of volatile compounds acting as semiochemicals, which influence the behavior of beetles as well as their predators (Davis, 2014) and modulate the growth of filamentous fungi, including mutualists of the beetle, entomopathogens, and opportunistic saprophytes (Hulcr & Dunn, 2011; Davis, 2014). Moreover, it has been suggested that yeasts provide nutritional supplements for the beetles; however, no direct evidence has yet supported this hypothesis (Six, 2013; Davis, 2014; Hofstetter *et al*., 2015).

The eastern larch beetle, *Dendroctonus simplex* LeConte, is a subcortical phloephagous insect that kills tamaracks, *Larix laricina* (Du Roi) K. Koch, and some exotic larch species. The distribution of this beetle extends throughout the range of the natural tree host, including northeastern and north-central of North America, western Canada, and Alaska (Langor & Raske, 1987a; Langor & Raske, 1987b; Seybold *et al*., 2002). This beetle is mainly considered as a secondary pest attacking freshly dead or weakened trees but, under epidemic conditions, it can also kill healthy *Larix* (Langor & Raske, 1987a; Langor & Raske, 1989). In the mid-1970s and early-1980s, a widespread outbreak of the eastern larch beetle caused the death of 1.4 million m^3^ of tamarack in the Atlantic provinces of Canada alone. Since then, the eastern larch beetle has been recognized as a serious forest pest (Langor & Raske, 1989). During the dispersal period, pioneer beetles attack trees and build galleries in the phloem layer. Following the reproduction and eggs hatching, larvae also excavate galleries and eat phloem throughout their development. The last larval instar digs a pupal chamber, stops feeding and empties his digestive tract in preparation for transformation in pupae, representing an inactive stage. Pupae will than transform into adults, overwintering until the next dispersal period (Langor & Raske, 1987a; Langor & Raske, 1987b). Although *D. simplex* is well studied, the associated fungi and their ecological roles in the development of this beetle remain to be investigated.

Since the relative importance of the filamentous fungi and yeasts associated with bark beetles likely varies over their different growth stages, the present study was undertaken to identify an overall portrait of fungal communities associated with *D. simplex* throughout its ontogeny, using 18S rRNA amplicon pyrosequencing. We compared fungal communities between different developmental stages of the larch beetle (*i.e*., adult, larva, and pupae). In a previous study, we have investigated the bacterial communities associated with the eastern larch beetle and found that the composition of bacterial communities is clearly dissimilar between the surface (ectomicrobiome) and the interior (endomicrobiome) of *D. simplex* body (Durand *et al*., 2015). Thus, these two microenvironments were also investigated in the present study. Additionally, the mycobiome associated with the galleries was also investigated. We hypothesize that the abundance of the associated fungi should vary according to the developmental stage of the beetle. These results would provide insights into the potential ecological roles of filamentous fungi and yeasts in the insect life cycle.

## Materials and Methods

### Site location, beetle processing, and samples preparation

Beetles were collected from a provincial larch plantation located near Saint-Claude (Quebec, Canada; Lat. 45.6809, Long. -71.9969) with the permission of the Ministère des Forêts, de la Faune et des Parcs authority. Log sections of randomly selected larch trees showing apparent signs of attacks by *D. simplex* were transported to the laboratory where they were stored at room temperature in plexiglass cages (30 cm x 30 cm x 88 cm). Beetle development was monitored weekly by gently peeling off the bark from the entry holes until the insects were reached. Based on their developmental morphology, pioneer beetles (adults), larvae, and pupae were randomly harvested with sterilized tweezers from different log sections, and insects were individually placed in sterile 2 ml microcentrifuge tubes.

For each developmental stage of *D. simplex*, the fungal microbiota associated with the ecto- and endomycobiome was recovered. For both of these fungal communities, three replicates were prepared following the method previously described (Durand *et al*., 2015). For each replicate, 50 insects were randomly selected, for a total of 150 insects per developmental stage. Briefly, for the ectomycobiome of each developmental stage, insects were pooled in 15 ml polypropylene tubes to recover sufficient fungal genomic DNA from the surface of the cuticle. Then, each sample underwent five serial washes with 5 ml phosphate-buffered saline (PBS) containing 0.1 % Triton X-100, with 1 min agitation (Genie 2 Vortex, Fisher, Ottawa, ON, Canada). The solution was filtered through a 0.22 μm nitrocellulose filter (EMD Millipore, Billerica, MA, USA) to recover the biomass. Each filter was placed in a Lysing matrix A tube (MP Biomedicals, Solon, OH, USA) for DNA extraction. To recover the endomycobiome from each developmental stage, ten previously washed beetles were randomly selected for each replicate. Their external surface was sterilized with three serial washes in 70% EtOH, followed by one wash with sterile water. The insects were then crushed into PBS and placed in a 2 ml screw cap tube containing 200 mg 0.1 mm glass beads (BioSpecs, Bartlesville, OK, USA) for DNA extraction.

The mycobiome associated with the subcortical galleries was recovered from the galleries where the pioneer beetles (adults) were collected. A total of 25 galleries were selected per replicate. First, insect frass was removed, and the inside galleries were carefully scraped using a sterile scalpel. For each selected gallery, the material was then placed in an individual sterile microtube. Samples were processed as for the ectomycobiome.

### DNA Extraction and PCR amplification

Total DNA was extracted following the method previously described (Durand *et al*., 2015). Briefly, 1 ml of extraction buffer containing 20 μg/ml RNase A was added to tubes containing the ecto- and endomycobiome. Cell lysis was achieved using the FastPrep^®^-24 Instrument (MP Biomedicals, Solon, OH, USA). Two cycles of lysis at 4 m/s for 50 s followed by 5 min on ice were performed consecutively. After centrifugation at 16,800 x *g* for 5 min, the supernatant was recovered, and extraction buffer containing RNase A was added to the previous tubes for the second cycle of lysis. Ammonium acetate was added to the supernatant at a final concentration of 2 M. The content was briefly mixed by inversion and the tubes kept on ice for 5 min before centrifugation at 20,800 x *g* for 15 min at 4°C. After collecting the supernatant, a second centrifugation was done with the same parameters. An equal volume of isopropyl alcohol (2-Propanol) was added to the supernatant, and DNA precipitation was performed overnight at 4°C. Centrifugation at 20,800 x *g* at 4°C for 30 min was done the next morning; then supernatant was discarded. Pellets were washed twice with 70% EtOH and were centrifuged at 20,800 x *g* for 15 min at 4°C. The EtOH supernatant was discarded, and pellets were air-dried before suspension in sterile ultrapure water. DNA concentration was estimated using the Quant-iT^™^ PicoGreen^®^ dsDNA Assay Kit (Invitrogen, Life Technologies, Burlington, ON, Canada) following the manufacturer instruction. The integrity of the genomic DNA was confirmed on a 1% agarose gel stained with ethidium bromide and visualized under UV light.

PCR amplification was achieved to confirm the presence of fungal DNA in the samples. Universal fungal primer NSA3 (5’ AAA CTC TGT CGT GCT GGG GAT A 3’) and NLC2 (5’ GAG CTG CAT TCC CAA ACA ACT C 3’) were used to amplified the SSU, ITS and LSU regions of the rRNA gene (Martin & Rygiewicz, 2005). Each 50 ul PCR reaction contained 25 mM MgCl_2_, 10 μg BSA, 10 mM dNTPs, 10 mM of each primer, 5 U Taq DNA polymerase and 10x ThermoPol^®^ buffer (New England Biolabs, Whitby, ON, Canada). Following the initial denaturation step of 5 min at 94°C, 30 amplification cycles were performed (94°C for 30 s, 67°C for 30 s, 72°C for 1 min) followed by a final extension step at 72°C for 10 min. Amplification was confirmed by electrophoresis of the PCR products on a 1.5% agarose gel stained with ethidium bromide and visualized under UV light. A negative control using all the extraction solutions but no insect was performed, and no amplification was observed.

### Fungal 18S rRNA pyrosequencing

DNA samples were sent to Research and Testing Laboratory (Lubbock, TX, USA) for sequencing. The fungal 18S rRNA gene was amplified using the universal primers SSUForward (5’ TGG AGG GCA AGT CTG GTG 3’) and funTitSsuRev (5’ TCG GCA TAG TTT ATG GTT AAG 3’). Roche 454 FLX-Titanium chemistry was used to sequence the amplicons. Elongation was performed from the forward primer. Raw data are available on NCBI under BioProject number PRJNA354793 for the developmental stages of D. simplex and PRJNA377102 for the galleries.

### Sequences processing pipeline

The post-sequencing processing were completed using the open-source program mothur v.1.33.0 software (<u>http://www.mothur.org</u>) (Schloss *et al*., 2009). Raw 454 reads were first processed to remove low quality reads, such as those containing (i) one or more uncertain bases (N), (ii) sequences shorter than 150 nt (nucleotides), (iii) unusually long reads that extended more than 100 nt over the amplicon size, (iv) reads that have long homopolymer sequences (more than 8), and (v) reads with incorrect forward primer sequences. Regions corresponding to the forward primer were kept to facilitate the alignment of the sequences during subsequent analyzes. Chimeras were removed with UCHIME against the SILVA reference alignment release 119 (Edgar *et al*., 2011; Quast *et al*., 2013), as implemented in mothur. The remaining filtered sequences were aligned by domain against the SILVA reference alignment using the ksize=9 parameter in mothur. Reads were also trimmed of all bases beyond the reverse primer with BioEdit 7.2.5 (<u>http://www.mbio.ncsu.edu/bioedit/bioedit.html</u>). Singletons were finally removed after clustering into draft Operational Taxonomic Units (OTUs) to obtain the final high quality reads. Libraries were normalized to the sequencing effort of the smallest 18S rRNA gene library (2641 sequences) to avoid biases in comparative analyzes introduced by the sampling depth. The final aligned reads were clustered into OTUs at ≥ 97% identity using the furthest neighboring cluster in mothur (Schmitt *et al*., 2012). Representative sequences of each OTU were taxonomically identified by BLASTN against the NCBI database (<u>http://www.ncbi.nlm.nih.gov</u>). The taxonomic identification is based on the current name in MycoBank Database (http://www.mycobank.org). Some contaminant OTUs related to protists were identified in a few samples; they were removed from the data set. The relative abundance of each OTU is presented.

### Diversity analysis

Rarefaction curves were generated using the mothur software. Diversity analyzes were performed with the R software 3.1.3 (<u>http://www.r-project.org</u>). Shannon index was calculated with the package “vegan”. ANOVA was performed with JMP Pro 12 (SAS Institute Inc., Cary, NC, USA) on obtained values. A Hellinger transformation was first applied to standardize the dataset. The “vegan” and “gclus” packages were used to generate the PCA. Equilibrium circle (not shown) of descriptor with the radius 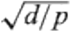 (where *d* is the number of dimensions of the reduced space: 2 and *p* is the total space: 1634) was plotted to identify OTUs significantly contributing to the axes defining the position of sampling sites. Additionally, a PERMANOVA was calculated with the package “vegan”. Indicator OTUs characterizing the different developmental stages were generated using the package “indicspecies”. Heatmap representing the distribution of these indicators across the developmental stages was generated using the “gplots” and “RColorBrewer” packages.

The representative sequences of each of the 69 OTUs found in high relative abundance throughout beetle developmental stages (≥1% of sample abundance), as well as their closely related sequences identified by BLASTN against NCBI database, were aligned together with the MUSCLE (Edgar, 2004) algorithm implemented in MEGA. A maximum likelihood phylogenetic tree was built with FastTree 2.1.7 (Price *et al*., 2010) using the GTR model with 1000 resampling to estimate node support values.

### Isolation and culture of *D. simplex* associated fungi

Yeasts and filamentous fungi associated with the adults of *D. simplex* were isolated using the same method previously described. The wash solutions and the crushed insects were plated on several culture media: Potato Dextrose Agar (pH 7.4; BD Difco, Franklin lakes, NJ, USA), Malt Extract Agar (pH 4.7; BD Difco), Czapek Solution Agar (pH 7.3; BD Difco) and Yeast Malt Extract Agar (pH 3.5; 0.3% yeast extract, 0.3% malt extract, 1% glucose, 0,5% tryptone, 2% agar). To only obtain fungi isolates, 1 mg/ml streptomycin, 0.05 mg/ml penicillin and 0.05 mg/ml chloramphenicol were added to the media. Petri dishes were incubated for two weeks at 25°C. Obtained fungi were purified separately on the same medium following the same conditions.

DNA extraction of the purified fungi was achieved using the same protocol as described above. A PCR amplification was done with the universal primers ITS1 (5’ TCC GTA GGT GAA CCT GCG G 3’) and ITS4 (5’ TCC TCC GCT TAT TGA TAT GC 3’) (White *et al*., 1990). After visualisation, PCR products were purified using the EZ-10 spin column PCR purification kit (Bio Basic, Markham, ON, Canada). A digestion with the restriction enzymes HaeIII, DdeI and HinfI (NEB) was done to select unique isolates. Selected purified PCR products were sequenced using the same primers and taxonomically identified by BLASTN. Isolates corresponding to OTUs identified by high-throughput sequencing are presented in supplemental data (Table S2).

## Results

### *Dendroctonus simplex* fungal diversity across developmental stages and microenvironments

To compare the fungal microbiota associated with the developmental stages and microenvironment of *D. simplex*, the ecto- and endomycobiome of the adults, larvae and pupae were recovered, using 50 insects for each of the three replicates per developmental stages. Following DNA extraction, a total of 18 samples were analysed, and 117,949 raw sequences were obtained. After quality control, 85,216 high-quality-filtered sequences were left. The average read length was 447 bp. After the equalization step and removing the protist reads, 44,377 sequences were kept for the remaining analyses. Clustering at 97% pairwise-identity threshold generated 1623 OTUs.

Rarefaction curves tend toward an asymptote, indicating that a suitable number of sequences were obtained for this analysis, although not all diversity was recovered (Fig S1). This indicates that a larger number of sequences will be needed in future studies. Additionally, no significant differences were observed in the diversity (Shannon index) between the analysed samples (Table S1).

A principal component analysis (PCA) was generated to see the dispersion of the samples along the axes of variation (Fig 1). Together, the first two axes explained 52.8% of the variation. The three replicates from all sample types clustered together, except one replicate associated with the endomycobiome of the adults, showing the evenness of the results. However, the dissimilarity of this particular sample did not influence the positions of the other samples in the PCA (data not shown). All replicates from the ectomycobiome of the adults and larvae are grouped together, showing similarity in the composition of their fungal communities. Samples belonging to the endomycobiome of larvae showed significant divergence from other samples, as they are separated by the second axis (19.3%). Finally, the pupae samples (ecto- and endo-) are separated from the two other developmental stages by the first axis (33.5%), showing a divergent OTUs assembly associated with this developmental stage.

**Figure 1.**
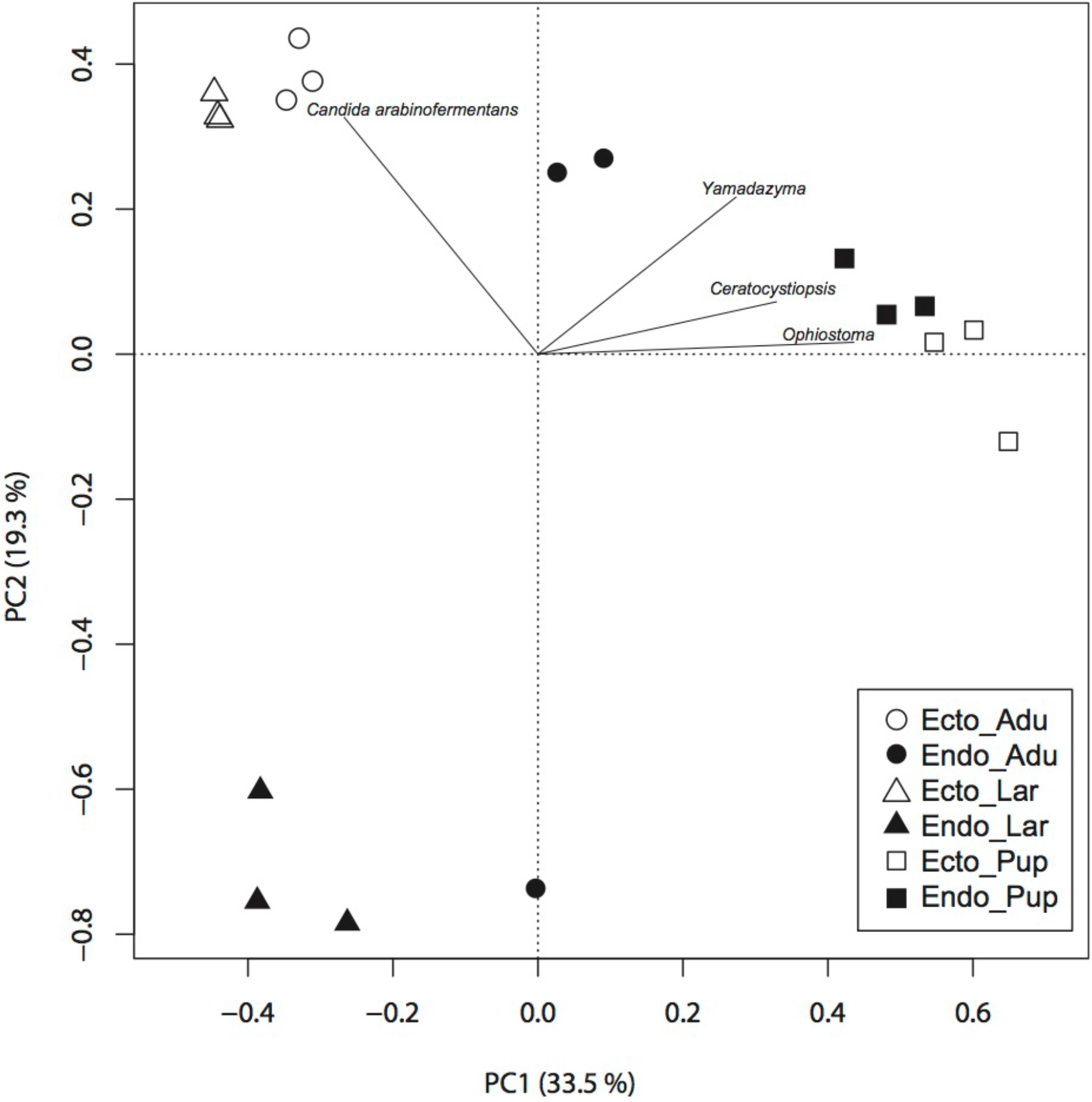
Principal component analysis of the different developmental stages and microenvironments of *D. simplex*. 50 insects were used per replicates, representing 150 insects for each developmental stage. All generated OTUs and equalized dataset were used to construct the PCA. Relative abundance of each OTU was used. “ECTO”, ectomycobiome; “ENDO”, endomycobiome; “ADU”, adults; “LAR”, larvae; “PUP”, pupae.

Four OTUs had a significant effect on the position of the samples in the PCA, represented by two yeasts and two filamentous fungi. *Candida arabinofermentans* influenced the position of the adults and larvae ectomycobiome, whereas one OTUs belonging to the *Yamadazyma* genus influenced the position of the adult’s endomycobiome. Two filamentous fungi genera, *Ceratocystiopsis* and *Ophiostoma*, influenced the position of pupae samples. Theses results seem to indicate that yeasts have an impact on the position of adults and larvae samples, whereas filamentous fungi influence the pupae samples position.

The PCA analysis seems to indicate that the developmental stages apply a selective pressure on the mycobiome, as shown by the position of the samples along the axes of variation. Accordingly, the developmental stage explained 59.1% of the variation observed in our analyses (PERMANOVA, p-value = 0.001).

### Taxonomic identification of mycobiome associated with developmental stages

Each of the 1623 OTUs linked to the developmental stages of the beetle was taxonomically identified by BLASTN against the NCBI database. A phylogenetic tree was generated with the abundant (≥ 1%) OTUs to confirm their taxonomic affiliations (Fig S2). Figure 2 shows the taxonomic identification of abundant (≥ 1% of sample relative abundance) yeast and filamentous fungi for each of the six microenvironments investigated. All identified yeasts and filamentous fungi belonged to the phylum Ascomycota. More specifically, all yeasts belong to the Saccharomycetes class, whereas filamentous fungi belonged to diverse classes. Yeasts were predominant in the adult ecto- and endomycobiomes, with more than 70% of the total abundance in each sample. Additionally, yeasts were also predominant in the larval ecto- and endomycobiome with, respectively, about 75% and 65% of the total abundance. On the other hand, filamentous fungi were most prevalent in the pupae, with 54% and 35% related, respectively, to ecto- and endomycobiomes. Depending on the samples, between 20 to 29% of the sequences observed were associated with non-abundant OTUs (<1% of the sample abundance). Among abundant OTUs, only one (1% to 20% according of the samples) was found in all microenvironments and was identified as *Candida oregonensis*. Additionally, one OTU assigned to the *Yamadazyma* genus (2% to 14%) was identified in all samples belonging to the ectomycobiome.

**Figure 2.**
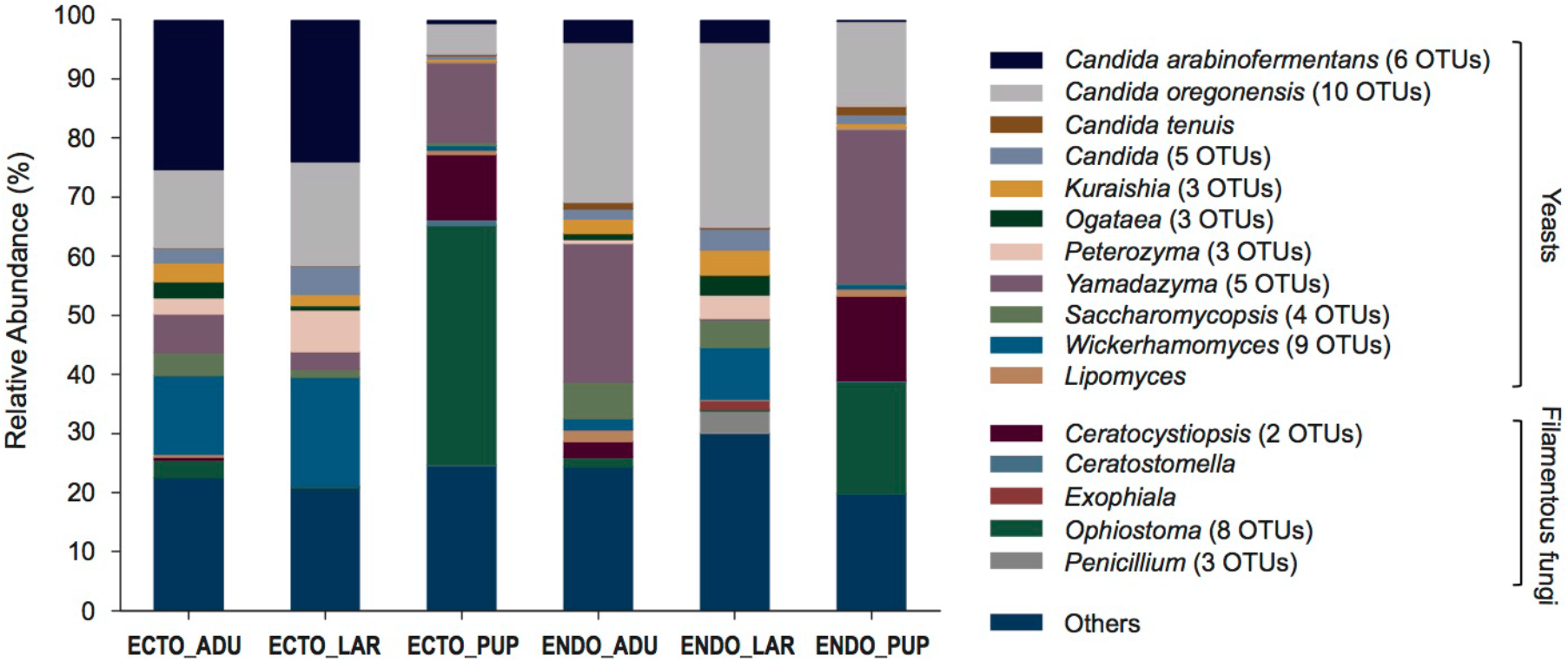
Taxonomical identification of the fungal community associated to the developmental stages and microenvironments of the eastern larch beetle. 50 insects were used per replicates, representing 150 insects for each developmental stage. All abundant OTUs (≥1% of the relative abundance per sample) are represented. Non-abundant OTUs (<1%) are grouped in the category others. Mean relative abundance of the three replicates is represented for each developmental stage. Yeasts identified to the gender represent cultured strains. “ECTO”, ectomycobiome; “ENDO”, endomycobiome; “ADU”, adults; “LAR”, larvae; “PUP”, pupae.

Among the identified yeasts, *Candida* species such as *C. arabinofermentans, C. tenuis* and *Candida* sp. were the most abundant, ranging from 5% to 30% of all OTUs. These OTUs were closely related to the ecto- and endomycobiome of adults and larvae, and under represented in pupae samples. On the other hand, OTUs corresponding to genera such as *Kuraishia, Ogataea and Peterozyma*, ranging from 1% to 7%, were also associated with both the ecto- and endomycobiome of adults and larvae, but not with the pupal samples. Additionally, *Saccharomycopsis* and *Wickerhamomyces* represented 1% to 13% of total abundance in adult and larval samples but were underrepresented in pupae samples. Finally, *C. oregonensis* and OTUs related to the *Yamadazyma* genus were found in all samples and were the two major yeast in pupal samples.

Among the identified filamentous fungi, *Ophiostoma* exhibited the highest abundance, with up to 40% in the pupae ectomycobiome. Moreover, theses OTUs were found mainly in the pupal samples but were also identified in the adult’s ecto- and endomycobiome, with abundance around 2%. *Ceratocystiopsis* genus was prevalent in pupal samples, with up to 14% of sample abundance. Additionally, this genus was also identified in the endomycobiome of the adults (3%). *Penicillium* was also found (3%) in the larvae endomycobiome, but was mostly absent in the other samples. Two others filamentous fungi genera were identified, such as *Ceratostomella* and *Exophiala*, with abundance close to 1%.

Indicator OTUs were calculated to define fungal OTUs that were closely associated with each developmental stage and microflora of *D. simplex* (Fig 3). Indicator OTUs associated with adult and larvae were all yeasts, whereas almost all OTUs associated with the pupae were filamentous fungi, reflecting the taxonomic identification (Fig 2). Several indicator OTUs were shared between the adults and larvae ectomycobiome, related to *C. arabinofermentans* and the genera *Peterozyma* and *Wickerhahomyces*, with higher abundance in larvae samples. Only one OTU, belonging to the *Ogataea* genus, was specific to the adult’s ectomycobiome. These results highlight, once again, the similarities between the ectomycobiome of adults and larvae. In contrast, indicator OTUs were specific for each developmental stage in the case of the endomycobiome of adults and larvae, with only one OTU abundantly present for the adults, identified as *C. oregonensis*. A greater number of indicator OTUs were identified for the larvae, mostly belonging to the genus *Candida*, and one OTU belonging to the genus *Ogataea*.

**Figure 3.**
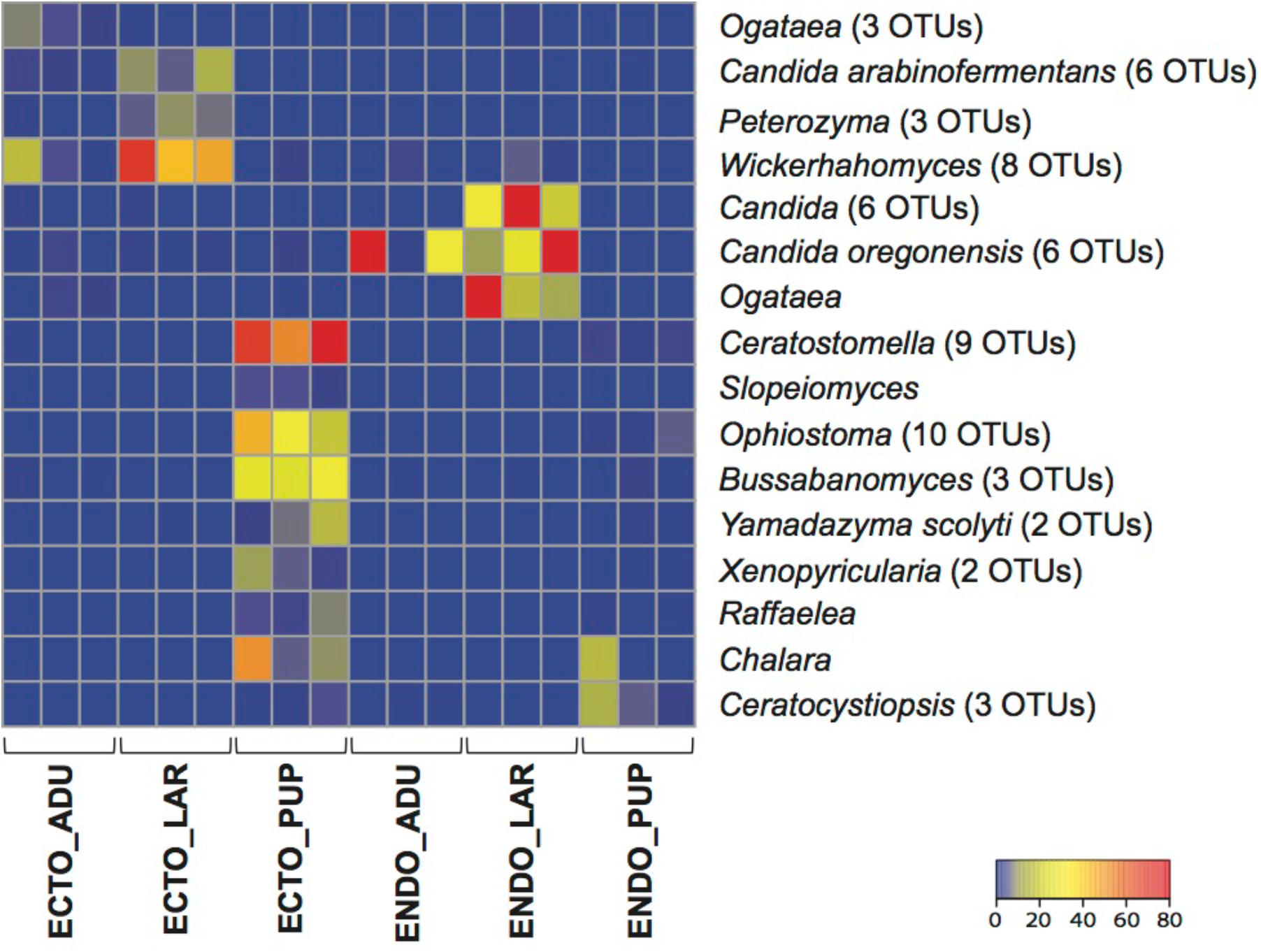
Indicator OTUs for each developmental stage and microenvironments of the eastern larch beetle. All generated OTUs and equalized dataset were used. All three replicates are shown on the heat map. Color key represent de number of sequences for the observed OTUs. Yeasts identified to the gender represent cultured strains. “ECTO”, ectomycobiome; “ENDO”, endomycobiome; “ADU”, adults; “LAR”, larvae; “PUP”, pupae.

Mostly filamentous fungi and one yeast were identified as indicator OTUs for both the ecto- and endomycobiome of the pupae. The ectomycobiome of the pupae is the niche where the larger number of specific indicator OTUs was found. Among them, *Ceratostomella, Ophiostoma* and *Yamadazyma* genera were identified. Other indicators, belonging to the genera *Slopeiomyces, Bussabanomyces, Xenopyricularia* and *Raffaelea* were also identified in lower abundance. Only two indicators OTUs were identified for the endomycobiome of pupae, belonging to the *Chalara* and *Ceratocystiopsis* genera. These results highlight, once again, the strong association of filamentous fungi with the pupal stage.

### Taxonomic identification of mycobiome associated with the environment of *D. simplex*

In order to identify the fungi diversity associated with the immediate environment of the beetle, the 438 OTUs associated with the galleries were taxonomically identified by BLASTN against NCBI database. Table 1 shows the identification of the abundant fungi for each replicate. All abundant OTUs were identified as yeast, such as *Candida* species that represent together 43% of the relative abundance. The *Wickerhamomyces* genus was also highly represented in the galleries (17%). Other genera of yeasts were identified with abundance ranging from 2% to 6%, such as *Yamadazyma, Ogatea, Kuraishia, Peterozyma* and *Saccharomycopsis*. All identified abundant OTUs associated with the galleries were also associated with the eastern larch beetle developmental stages (Fig 2). Additionally, some non-abundant OTUs were found, representing 19% of samples relative abundance.

**Table 1.** Taxonomical identification of the fungal community associated to the environment of the eastern larch beetle. All abundant OTUs (≥1% of the relative abundance per sample) are represented in the table, non-abundant OTUs (<1%) are grouped in the category others. The mean relative abundance of the three replicates is presented.

| Taxonomical identification | Relative abundance |
| --- | --- |
| *Candida* (9 OTUs) | 43.77 |
| *Yamadazyma* (2 OTUs) | 3.96 |
| *Ogataea* (4 OTUs) | 6.13 |
| *Kuraishia* (2 OTUs) | 3.85 |
| *Peterozyma* (2 OTUs) | 3.75 |
| *Saccharomycopsis* (2 OTUs) | 1.67 |
| *Wickerhamomyces* (6 OTUs) | 17.23 |
| Others | 19.06 |

## Discussion

The aim of this work was to characterize the mycobiome (filamentous fungi and yeasts) associated with the different developmental stages of the eastern larch beetle. Associations between several *Dendroctonus* species and their fungal partners have been widely reviewed (Popa *et al*., 2012; Six, 2013). However, the mycobiome of the eastern larch beetle has never been comprehensively studied. Here, we investigated the fungal diversity associated with the ecto- and endomycobiomes of adult, larvae, and pupae developmental stages of *D. simplex* and their immediate environment using 18S rRNA pyrosequencing. Our results showed that yeasts were mainly associated with the adults and larvae, whereas filamentous fungi were prevalent in the pupal stage.

Great fungi diversity was identified for each developmental stage and microflora. The fact that fungi are present throughout the ontogeny of the beetle suggests that they are probably essential for the insect’s development. No statistically significant differences were observed between the ecto- and endomycobiome of all developmental stages in term of diversity, which indicates that fungal flora is equally distributed at the surface of the cuticle and in the interior of the insect. A variety of filamentous fungi and yeasts can accomplish a single essential function within the beetle microbiome (Hofstetter *et al*., 2015). The eastern larch beetle could then benefit from transporting a wide variety of associated fungi, promoting the adaptation to a broader range of environments. Although Davis (2014) mentioned that bark beetles harbour a lower yeast diversity compared to other families of beetles, we found a significant yeast diversity throughout the developmental stages of *D. simplex*. In fact, yeasts were more abundant than filamentous fungi (50 OTUs and 19 OTUs found in abundance respectively). Filamentous fungi associated with bark beetles have been reported to interfere with the tree defense systems, as well as other possible function, impeding tree defenses (Paine *et al*., 1997; Six & Wingfield, 2011). As the eastern larch beetle attacks mostly weakened or freshly dead trees, the presence of filamentous fungi may be less critical, allowing a greater proportion for insect-associated yeasts. The higher diversity observed in this study compared to previous ones may also be due to a bias associated with the technique used. Most previous studies on bark beetles employed culture-dependant or molecular methods (Adams & Six, 2007; Khadempour *et al*., 2012; Lou *et al*., 2014; Hu *et al*., 2015), whereas here we used high throughput sequencing, which allowed the identification of the majority of the specific richness, explaining the higher number of observed species.

Fungal diversity associated with the ectomycobiome of adults and larvae is similar to each other, but distinct from that of the pupal stage, as shown by the PCA and the higher relative abundance of yeasts in the adults and larvae stages. This shift in fungal communities can be explained by the fact that pupae are an inactive stage found in a closed environment, and that they do not feed. Oppositely, adults and larvae share the same environment under the bark. During the construction of galleries, mainly yeasts, and filamentous fungi, carried by the adults could be inoculated into the galleries. Accordingly, 10 species of yeasts identified in the ectomycobiome of adults and larvae were also identified in the beetle’s galleries, which represent 80% of galleries abundance. Oppositely, only one species of filamentous fungi found in the ectomycobiome of the adults was identified in the galleries. Yeasts are frequently isolated from adults and larvae of bark beetles (Davis, 2014; Hofstetter *et al*., 2015). Almost all the identified yeasts in this study were also previously recovered from various beetles (Davis, 2014). The endomycobiome of the adults and larvae, even if they represent two distinct populations as shown on the PCA, are also composed of a great majority of yeast OTUs. Only a few filamentous fungi were found in these microenvironments. During the beetle development, adults and larvae feed on bark as they construct their galleries (Langor & Raske, 1987b; Wood, 2007). Simultaneously, they feed on yeasts and filamentous fungi found in their galleries, which significantly reduced their development period (Paine *et al*., 1997; Six & Paine, 1998; Klepzig & Six, 2004; Davis, 2014). However, even if the nutritional benefits of yeasts have often been proposed for bark beetles, no study clearly demonstrates this phenomenon (Davis, 2014). The high prevalence of yeasts associated with the adults and larvae and the presence of small proportions of filamentous fungi associated with their endomycobiomes are supporting the hypothesis that yeasts are essential nutritional elements for the development of the eastern larch beetle and that the insect is performing some microbial gardening. Some insects, such as ants, termites and ambrosia beetles, transport and cultivate their fungal associates in order to feed on them (Mueller & Gerardo, 2002; Mueller *et al*., 2005). Furthermore, fungus gardens of scolytine ambrosia beetles are composed of an assemblage of filamentous fungi, yeasts and bacteria (Mueller *et al*., 2005). This fungus-farming behaviour has never been observed in bark beetles to date. This type of behaviour, *i.e*. inoculating their fungal associates in galleries in order to feed on them, could explain the similarities observed between the adults and larvae mycobiome. Yeasts and filamentous fungi associated with *Dendroctonus* species provide the insect with essential nutriments, such as sterols, reducing the developmental time of larvae and increasing beetle fitness (Six & Paine, 1998; Bentz & Six, 2006). Moreover, yeasts such as *Candida* species can assimilate nutrient such as nitrate, xylose and cellobiose (Lou *et al*., 2014). Nutritional need should be different over the developmental stages of the beetle, as the beetle needs nutritional benefits to accomplish different steps in is life cycle, such as ovogenesis and development. These differences in requirements could reflect the greater distance observed associated with the endomycobiome, whereas different yeasts could provide the insect with different nutrients, supporting once again the nutritional hypothesis for yeasts.

In the literature, many functions have been associated with yeasts other than nutrition, such as mediation of beetle-microorganism interactions, production of volatiles compounds or degradation of tree chemicals (Davis, 2014). Theses function can be accomplished by many genera of yeasts, while a single yeast may accomplish many functions (Six, 2013; Davis, 2014). Pioneer beetles were collected for this study. During the beetle attack, the trees release volatile compounds, such as terpenes, that are toxic to the insects (Adams *et al*., 2011; Davis & Hofstetter, 2012). Some yeast associated with the eastern larch beetle may detoxify the environment, facilitating the adaptation of the insect. It has been shown that *O. pini*, associated to female *D. brevicomis*, can alter concentration of monoterpenes present in phloem tissue of *Pinus ponderosa* over time (Davis & Hofstetter, 2011). Accordingly, *Ogataea* was found in abundance in the adults and larvae ectomycobiome.

A few filamentous fungi were identified in samples coming from the ecto- and endomycobiome of the adults and larvae, such as *Ophiostoma* and *Ceratocystiopsis*. These genera are grouped as ophiostomatoids fungi, which are commonly associated with bark beetles (Paine *et al*., 1997; Six & Wingfield, 2011; Six, 2013; Hofstetter *et al*., 2015). Theses fungi help beetles to overcome tree defenses by colonizing resin ducts, in order to proceed to a successful attack (Paine *et al*., 1997). This function may explain the presence of a few ophiostomatoids fungi associated mainly with the adult’s samples, responsible for the initial attack and detoxification of the immediate environment. Among other roles, nutrition and mediation of microbial associations have been proposed (Paine *et al*., 1997).

The mycobiomes associated with the surface and the interior of the pupae are separated from the other developmental stages of the beetle, as shown on the PCA. Under the bark, pupae are found in a pupal chamber closed by frass, forming a restricted environment (Langor & Raske, 1987a). This microenvironment likely explains why pupae have such a different mycobiome from the other two developmental stages, especially represented by a majority of filamentous fungi. Indeed, indicator OTUs for this developmental stage are almost all filamentous fungi, showing a strong association between them. For instance, *Ophiostoma* was more abundant in these samples (41% in the ectomycobiome and 19% in the endomycobiome) than in the adult’s samples (3% and 2% respectively). Others ophiostomatoids fungi were also identified in those samples. During the pupal stages, beetles stop feeding and undergo a reorganization of their body (Langor & Raske, 1987a). This finding could explain why yeast abundance was less important in these samples, once again supporting the hypothesis of a nutritional function for these associates. Pupae are also more fragile than the other developmental stages of the beetle. Filamentous fungi associated with the bark beetles can protect the insect against antagonistic fungi by inhibiting their growth (Paine *et al*., 1997). Pupae may need these filamentous fungi for protection during this crucial period of development.

Important fungal diversity was identified associated with developmental stages of the eastern larch beetle. Yeasts were more abundant than filamentous fungi in the samples coming from the adults and larvae, presumably indicating a nutritional benefit from this type of microorganism. In contrast, fungi were more abundant at the pupae stages, possibly reflecting a role in protection. Further studies are required to confirm these functions. Our results reveal that the eastern larch beetle is closely associated with various yeasts and filamentous fungi throughout its development cycle, with varying relative abundances according to the developmental stages. We propose that microbial gardening behaviour is probably involved as yeasts and filamentous fungi seem to play essential roles in the development of the eastern larch beetle. Bacteria and fungi are important partners in the *D. simplex* microbiome. Now that the bacterial and fungal microbiomes of this beetle have been characterized, it will be interesting to study the ecological interactions between these two types of microorganisms, in order to better understand the mechanism of *D. simplex* attack and development under the bark.

## Funding

This work was supported by the Direction Générale de la Production des Semences et de Plants Forestiers (DGPSP) [grant number DGPSP-2013-1122435] to CG. AAD was supported by a Wladimir A. Smirnoff Fellowship.

## Acknowledgments

We would like to thank Amélie Bergeron for her help with samples preparation and DNA extraction. We would also like to thank Fabrice Jean-Pierre and Marie-Christine Groleau for technical assistance.

## References

Adams AS, Aylward FO, Adams SM, Erbilgin N, Aukema BH, Currie CR, Suen G & Raffa KF (2013) Mountain pine beetles colonizing historical and naïve host trees are associated with a bacterial community highly enriched in genes contributing to terpene metabolism. Appl. Environ. Microbiol. 79: 3468–3475.

Adams AS, Boone CK, Bohlmann J & Raffa KF (2011) Responses of bark beetle-associated bacteria to host monoterpenes and their relationship to insect life histories. J. Chem. Ecol. 37: 808–817.

Adams AS & Six DL (2007) Temporal variation in mycophagy and prevalence of fungi associated with developmental stages of *Dendroctonus ponderosae* (Coleoptera: Curculionidae). Environ. Entomol. 36: 64–72.

Ayres MP, Wilkens RT, Ruel JJ, Lombardero MJ & Vallery E (2000) Nitrogen budgets of phloem-feeding bark beetles with and without symbiotic fungi. Ecology 81: 2198–2210.

Bentz BJ & Six DL (2006) Ergosterol content of fungi associated with *Dendroctonus ponderosae* and *Dendroctonus rufipennis* (Coleoptera: Curculionidae, Scolytinae). Ann. Entomol. Soc. Am. 99: 189–194.

Bleiker KP & Six DL (2007) Dietary benefits of fungal associates to an eruptive herbivore: Potential implications of multiple associates on host population dynamics. Environ. Entomol. 36: 1384–1396.

Davis TS (2014) The Ecology of Yeasts in the Bark Beetle Holobiont: A Century of Research Revisited. Microb. Ecol. 69: 723–732.

Davis TS & Hofstetter RW (2011) Reciprocal interactions between the bark beetle-associated yeast *Ogataea pini* and host plant phytochemistry. Mycologia 103: 1201–1207.

Davis TS & Hofstetter RW (2012) Plant secondary chemistry mediates the performance of a nutritional symbiont associated with a tree-killing herbivore. Ecology 93: 421–429.

Durand A-A, Bergeron A, Constant P, Buffet J-P, Déziel E & Guertin C (2015) Surveying the endomicrobiome and ectomicrobiome of bark beetles: The case of *Dendroctonus simplex*. Sci Rep 5: 17190.

Edgar RC (2004) MUSCLE: multiple sequence alignment with high accuracy and high throughput. Nucleic Acids Res. 32: 1792–1797.

Edgar RC, Haas BJ, Clemente JC, Quince C & Knight R (2011) UCHIME improves sensitivity and speed of chimera detection. Bioinformatics 27: 2194–2200.

Hofstetter RW, Dinkins-Bookwalter J, Davis TS & Klepzig KD (2015) Symbiotic associations of bark beetles. Bark Beetles, Vega F E & Hofstetter R W (Édit.) Elsevier, USA.

Hu X, Li M & Chen H (2015) Community structure of gut fungi during different developmental stages of the Chinese white pine beetle (*Dendroctonus armandi*). Sci Rep 5.

Hulcr J & Dunn RR (2011) The sudden emergence of pathogenicity in insect-fungus symbioses threatens naive forest ecosystems. Proceedings of the Royal Society B: Biological Sciences 278: 2866–2873.

Khadempour L, LeMay V, Jack D, Bohlmann J & Breuil C (2012) The relative abundance of mountain pine beetle fungal associates through the beetle life cycle in pine trees. Microb. Ecol. 64: 909–917.

Klepzig KD & Six DL (2004) Bark beetle-fungal symbiosis: Context dependency in complex associations. Symbiosis 37: 189–205.

Langor DW & Raske AG (1987a) Emergence, host attack, and overwintering behavior of the eastern larch beetle, *Dendroctonus simplex* LeConte (Coleoptera: Scolytidae), in Newfoundland. The Canadian Entomologist 119: 975–983.

Langor DW & Raske AG (1987b) Reproduction and development of the eastern larch beetle, *Dendroctonus simplex* LeConte (Coleoptera: Scolytidae), in Newfoundland. The Canadian Entomologist 119: 985–992.

Langor DW & Raske AG (1989) The eastern larch beetle, another threat to our forests (Coleoptera: Scolytidae). Forestry Chronicle 65: 276–279.

Lou Q-Z, Lu M & Sun J-H (2014) Yeast diversity associated with invasive *Dendroctonus valens* killing *Pinus tabuliformis* in China using culturing and molecular methods. Microb. Ecol.

Martin KJ & Rygiewicz PT (2005) Fungal-specific PCR primers developed for analysis of the ITS region of environmental DNA extracts. BMC Microbiol. 5: 1–11.

Moore ML & Six DL (2015) Effects of temperature on growth, sporulation, and competition of mountain pine beetle fungal symbionts. Fungal Microbiology 70: 336–347.

Mueller UG & Gerardo NM (2002) Fungus-farming insects: Multiple origins and diverse evolutionary histories. PNAS 99: 15247–15249.

Mueller UG, Gerardo NM, Aanen DK, Six DL & Schultz TR (2005) The evolution of agriculture in insects. Annual Review of Ecology, Evolution, and Systematics 36: 563–595.

Paine TD, Raffa KF & Harrington TC (1997) Interactions among scolytid bark-beetles, their associated fungi, and live host conifers. Annu. Rev. Entomol.:179–206.

Popa V, Déziel E, Lavallée R, Bauce E & Guertin C (2012) The complex symbiotic relationships of bark beetles with microorganisms: A potential practical approach for biological control in forestry. Pest Manag. Sci. 68: 963–975.

Price MN, Dehal PS & Arkin AP (2010) FastTree 2 – Approximately maximum-likelihood trees for large alignments. PLoS ONE 5: e9490.

Quast C, Pruesse E, Yilmaz P, Gerken J, Schweer T, Yarza P, Peplies J & Glöckner FO (2013) The SILVA ribosomal RNA gene database project: improved data processing and web-based tools. Nucleic Acids Res. 41: D590–D596.

Rivera FN, Gonzalez E, Gomez Z, Lopez N, Hernandez-Rodriguez C, Berkov A & Zuniga G (2009) Gut-associated yeast in bark beetles of the genus *Dendroctonus* Erichson (Coleoptera: Curculionidae: Scolytinae). Biol. J. Linn. Soc. 98: 325–342.

Schloss PD, Westcott SL, Ryabin T, Hall JR, Hartmann M, Hollister EB, Lesniewski RA, Oakley BB, Parks DH, Robinson CJ, Sahl JW, Stres B, Thallinger GG, Van Horn DJ & Weber CF (2009) Introducing mothur: open-source, plaform-independent, community-supported software for describing and comparing microbial communities. Applied and Environmental Microbiology 75: 7537–7541.

Schmitt S, Tsai P, Bell J, Fromont J, Ilan M, Lindquist N, Perez T, Rodrido A, Schupp PJ, Vacelet J, Webster NS, Hentschel U & Taylor MW (2012) Assessing the complex sponge microbiota: core, variable and species-specific bacterial communities in marine sponges. The ISME Journal 6: 564–576.

Seybold SJ, Albers MA & Katovich SA (2002) Eastern Larch Bettle. Forest Insects & Disease Leaflet 175: 1–10.

Shi W, Guo Y, Xu C, Tan S, Miao J, Feng Y, Zhao H, St. Leger RJ & Fang W (2014) Unveiling the mechanism by which microsporidian parasites prevent locust swarm behavior. PNAS 111: 1343–1348.

Six DL (2003) A comparation of mycangial and foretic fungi of individual mountain pine beetles. Can. J. For. Res. 33: 1331–1334.

Six DL (2012) Ecological and evolutionary determinants of bark beetle — fungus symbioses. Insects 3: 339–366.

Six DL (2013) The bark beetle holobiont: Why microbes matter. J. Chem. Ecol. 39: 989–1002.

Six DL, De Beer ZW, Duong TA, Carroll AL & Wingfield MJ (2011) Fungal associates of the lodgepole pine beetle, *Dendroctonus murrayanae*. Antonie van Leeuwenhoek, International Journal of General and Molecular Microbiology 100: 231–244.

Six DL & Klepzig KD (2004) Dendroctonus Bark Beetles as Model Systems for Studies on Symbiosis. Symbiosis 37: 207–232.

Six DL & Paine TD (1998) Effects of mycangial fungi and host tree species on progeny survival and emergence of *Dendroctonus ponderosae* (Coleoptera: Scolytidae). Environ. Entomol. 27: 1393–1401.

Six DL & Wingfield MJ (2011) The role of phytopathogenicity in bark beetle-fungus symbioses: a challenge to the Classic Paradigm. Annu. Rev. Entomol.:255–272.

White TJ, Bruns T, Lee S & Taylor J (1990) Amplification of direct sequencing of fungal ribosomal RNA genes for phylogenetics. Academic Press, New York.

Wood SL (2007) Bark and ambroza beetles of South America (Coleoptera: Scolytidae). Brigham Young University, Provo, Utah USA. 900 p.

